# Antigenic Sensitivity of Membrane-Proximal Targeting Chimeric Antigen Receptors can be Fine-Tuned through Hinge Truncation

**DOI:** 10.1101/2020.10.30.360925

**Authors:** Scott McComb, Tina Nguyen, Alex Shepherd, Kevin A. Henry, Darin Bloemberg, Anne Marcil, Susanne Maclean, Rénald Gilbert, Christine Gadoury, Rob Pon, Traian Sulea, Qin Zhu, Risini D. Weeratna

## Abstract

**Background:** Chimeric antigen receptor (CAR) technology has revolutionized the treatment of B-cell malignancies and steady progress is being made towards CAR-immunotherapies for solid tumours. Epidermal growth factor family receptors EGFR or HER2 are commonly overexpressed in cancer and represent proven targets for CAR-T therapy; given their expression in healthy tissues it is imperative that any targeting strategy consider the potential for on-target off-tumour toxicity.

**Methods:** Herein, we utilize high-throughput CAR screening to identify novel camelid single-domain antibody CARs (sdCARs) with high EGFR-specific CAR-T response. To optimize antigenic sensitivity of this EGFR-sdCAR, we performed progressive N-terminal truncation of the human CD8 hinge domain used as a spacer in many CAR constructs. Hinge truncation resulted in decreased CAR sensitivity to EGFR and improved selectivity for EGFR-overexpressing cells over EGFR-low target cells or healthy donor derived EGFR-positive fibroblasts. To investigate the molecular mechanism of hinge truncation, we test hinge-truncated scFv-based CARs targeting membrane proximal or membrane distal domains of EGFR-family proteins, HER2 and EGFRvIII. Finally, we proceed to test hinge variant EGFR-sdCAR functionality through *in vitro* and *in vivo* assessments in primary T cells derived from multiple donors.

**Results:** For CARs targeting membrane-proximal epitopes, hinge truncation by even a single amino acid provided fine control of the antigenic sensitivity, whereas CARs targeting membrane distal domains were not sensitive to even complete hinge domain removal. Hinge-modified EGFR-sdCARs showed consistent and predictable responses in Jurkat-CAR cells and primary human CAR-T cells *in vitro* and *in vivo*.

**Conclusions:** Overall, these results indicate that membrane-proximal epitope targeting CARs can be modified through hinge length tuning for programmable antigenic sensitivity and improved tumour selectivity.

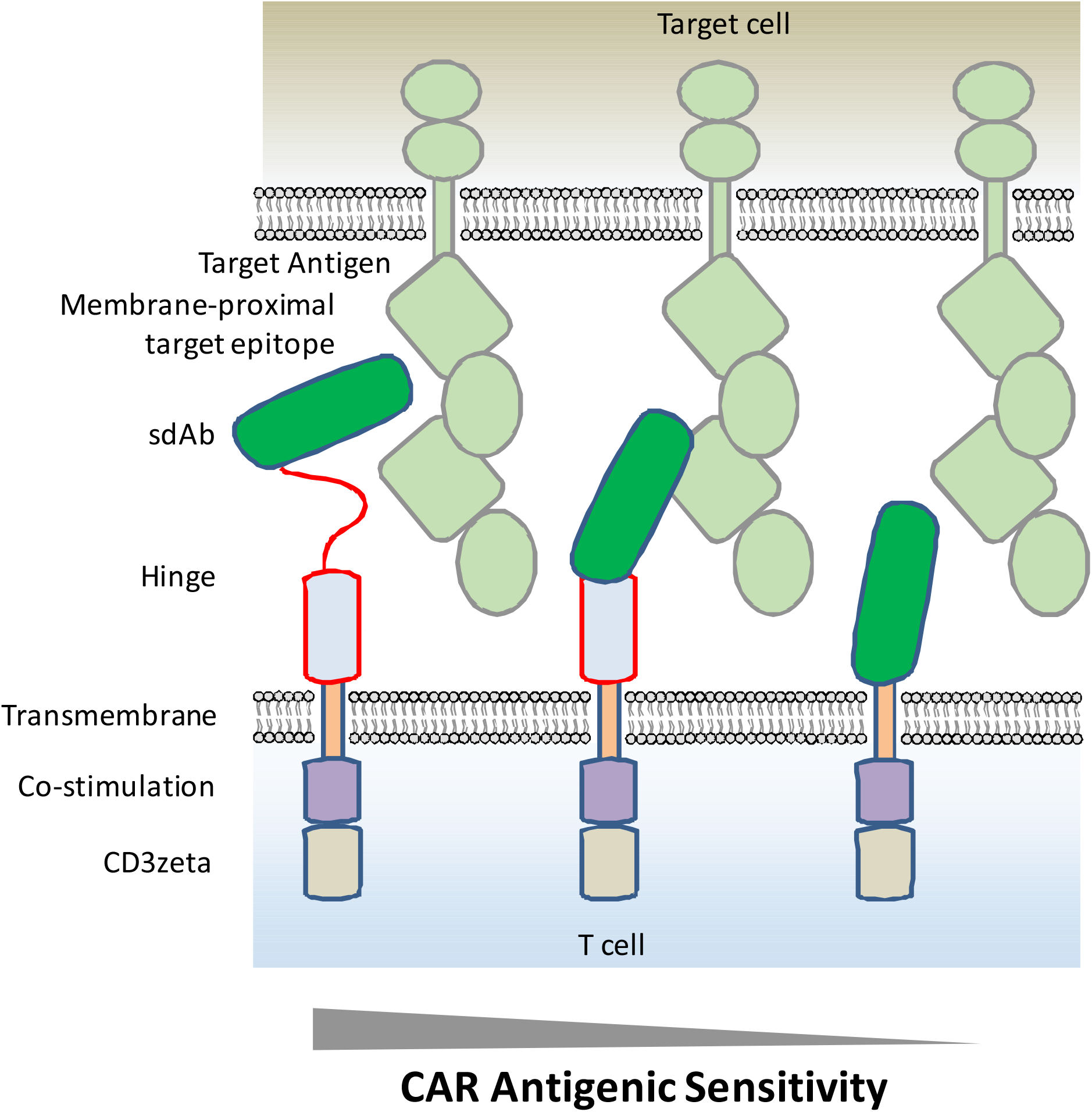

- Single amino acid truncations of CD8-hinge domain provide precise control of CAR antigen sensitivity
- Truncated hinge CARs show enhanced selectivity for antigen overexpressing tumour cells and decreased activity towards healthy antigen-expressing cells
- Epitope location is a critical factor in determining hinge sensitivity for a CAR
- Hinge tuning can modulate CAR-T antigenic sensivity *in vitro* and *in vivo*

## Background

Following the remarkable clinical success of CD19-targeted chimeric antigen receptor (CAR) therapies for the treatment of B-cell malignancies, design and development of novel CARs for more common solid tumours is a highly active area of research and development[1]. The first step for CAR-development is the identification of an antigen binding domain (ABD), typically composed of an antibody single-chain variable fragment (scFv)[2,3], which can be used to create a CAR with high antigen-specific responsive activity. Previous work has also demonstrated that camelid single-domain antibodies (sdAb), also called V_H_Hs or nanobodies, can be also be used as effective CAR ABDs[4,5], and an sdAb-based CARs is now producing promising results as a clinical CAR-T therapy for myeloma[6,7]. After identification of a functional ABD, molecular optimization can be used to further fine-tune the therapeutic properties of a CAR through alteration of various elements of the CAR molecule including: the ABD, hinge, transmembrane domain, and intracellular signaling domains; each of which have been demonstrated to have significant impact on the signaling properties and functionality of CARs[2,3].

Recent work has shown that decreased CAR signaling in CD19-specific CARs can be achieved by lowering the affinity of the ABD[8], altering the hinge or transmembrane domains[9–11], or engineering signaling domains with reduced activity[12]. Intriguingly, these diverse molecular strategies have enhanced CAR-T persistence and therapeutic benefit in animal models and in at least one clinical trial thus far[8], suggesting that an optimal level of signaling may be somewhat lower than that of current generation clinical CD19-targeting CAR-T therapies. Thus, molecular optimization of CARs has emerged as a viable strategy for widening the therapeutic window of novel CAR-T therapies.

Epidermal growth factor receptor (EGFR) is one of the most commonly altered oncogenes in solid cancers, either through a variety of activating mutations or through over-expression of the native receptor[13,14]. EGFR has a relatively large extracellular domain with four subdomains[15], and is a well-established target for monoclonal antibodies and small-molecule inhibitors[16,17]. EGFR has also been explored as a target for CAR-T therapy, and clinical trials have been undertaken using ABDs specific for either the WT[18,19] or a mutated tumour-specific form of the receptor known as EGFRvIII[20,21]. Recent clinical reports using EGFR-targeted CAR-T therapy in lung, biliary, and pancreatic cancers revealed no unmanageable toxicity and documented partial disease responses in some patients[22–25]. EGFR is also under investigation as a target for bi-specific immune engaging therapy[26].

As EGFR is expressed in normal tissues as well as on tumour cells it is imperative that any targeting strategy consider the potential for on-target off-tumour toxicity. In pre-clinical CAR-T work, it has previously been shown that the use of lower affinity EGFR or human epidermal growth factor 2 (HER2) ABDs for CAR-T can improve selectivity for overexpressing tumour cells over normal tissues[27]. Herein we present novel camelid single-domain antibody (sdAb) CAR molecules with high on-target activity against human EGFR (EGFR-sdCAR), and demonstrate that truncation of the hinge domain can be used to fine-tune CAR antigenic sensitivity and enhance selectivity for antigen overexpressing tumours. Similar hinge truncation with scFv-CARs targeting HER2 and EGFRvIII support a view the hinge/spacer CARs is critical only for CARs targeting membrane-proximal epitopes. Thus, our results indicate that hinge truncation offers a powerful and precise means to customize antigenic sensitivity of CARs targeting membrane proximal domains of EGFR or other cancer associated antigens, and could be a useful tool to better understanding CAR biology and for the development of more complex multi-antigen targeting CAR therapeutics.

## Methods

### CAR cloning

Three previously reported EGFR-specific sdAb sequences[28] were cloned into a modular CAR backbone using PCR amplification and single-pot restriction ligation as previously described[29]. EGFR-sdCAR constructs bearing either a full-length human 45 amino acid CD8 hinge (45CD8h) or progressively N-terminally truncated hinge variants (34CD8h, 22CD8h, or no hinge) were cloned using Gibson assembly. A library of sdAb021-CAR truncation mutants with single amino acid N terminal truncations of the human CD8 hinge extended with an additional N-terminal flexible linker [(GGGGS)_3_GG-CD8h] was generated using a modular hinge-CAR with convenient type-IIs restriction sites integrated into the construct 3’ of the sdAb coding region. An array of DNA encoding truncated CD8 hinge domains of varying lengths (all possible variants between 60 and 1 amino acid) were synthesized as DNA fragments (Twist Bioscience, USA) and cloned into the sdAb021-modular-hinge-BBz-GFP CAR construct using single-pot restriction ligation. Limited hinge truncation libraries with defined-target CARs were generated by exchanging the sdAb021 sequence with HER2 or EGFRvIII specific scFv sequences. Trastuzumab derived scFv sequences were generated based on previously reported mutant forms of trastuzumab with enhanced avidity[30], whereas EGFRvIII-targeting scFvs were generated as previously reported[29]. Both HER2- and EGFRvIII-scFvs were in a VH-(G4S)_3_-VL format.

### Cell lines and culture

Other than U87MG and HDF cells, all others were obtained from American Tissue Culture Collection (ATCC, Manassas, VA). The glioblastoma cell line U87 MG-WT and U87 MG-vIII (U87-vIII, expressing EGFRvIII via retroviral transduction and sorting) were kindly provided by Professor Cavnee, from the Ludwig Institute for Cancer Research, University of California, San Diego (San Diego, CA, USA)[31]. The human dermal fibroblasts (HDF) were purchased from Cell Applications, Inc (San Diego, CA, USA). The T-cell lines used were Jurkat, and target cells SKOV3, MCF7, U87 MG vIII, Raji, and Nalm6. Target cells were transduced with lentivirus containing NucLight Red (Sartorius, Essen BioScience, USA), a third generation HIV-based, VSV-G pseudotyped lentivirus encoding a nuclear-localized mKate2. Nuclight positive cells were obtained by selection with puromycin. HDF were cultured in all in one ready to use fibroblast growth medium (Cell Applications, Inc). U87 MG, SKOV3, MCF7, were cultured in DMEM supplemented with 10% fetal bovine serum (FBS), 2mM L-glutamine, 1mM sodium pyruvate and 100 µg/mL penicillin/streptomycin. Jurkat, Raji, and Nalm6 were cultured in RPMI supplemented with 10% FBS, 2mM L-glutamine, 1mM sodium pyruvate and 100 µg/mL penicillin/streptomycin. These cell lines were tested for the presence of mycoplasma contamination by PCR.

### CAR-J assay

High-throughput assessments of CAR function were performed by CAR-J, according to previously outlined protocol[29] . Briefly, 5×10^5 cells were suspended in 100 µl Buffer 1SM (5 mM KCl, 15 mM MgCl_2_, 120 mM Na_2_HPO_4_/NaH_2_PO_4_, 25 mM sodium succinate, and 25 mM mannitol; pH7.2) and incubated with 2µg of pSLCAR-CAR plasmids as described in the text or with no plasmid control. Cells and plasmid DNA in solution were transferred into 0.2 cm generic electroporation cuvettes (Biorad Gene Pulser; Bio-Rad Laboratories, Hercules, California, USA) and immediately electroporated using a Lonza Nucleofector I (Lonza, Basel, Switzerland) and program X-05 (X-005 on newer Nucleofector models). Cells were cultured in pre-warmed recovery media (RPMI containing 20% FBS, 1mM sodium pyruvate and 2 mM L-glutamine) for four 4h before being co-cultured with EGFR-expressing target cells U87 MG-vIII, MCF7 and SKOV-3 or negative control Ramos and Nalm6. Electroporated Jurkat cells were added to varying numbers of target cells in round bottom 96-well plates in effector to target (E:T) ratios ranging from 1:10 to 100:1 (effector to target ratio) or with no target cells (or an E:T of 1:0) and cultured overnight before being staining with allophycocyanin (APC)-conjugated anti human-CD69 antibody (BD Biosciences #555533). Flow cytometry was performed using a BD- Fortessa (BD Biosciences) and data was analyzed using FlowJo software (FlowJo LLC, Ashland, Oregon, USA) and visualized using GraphPad Prism (GraphPad Software, Inc. California, USA).

### Plate-bound anti-CD3 CAR-J Stimulation Assay

Stable Jurkat cells expressing hinge variant sdAb021-BBz CARs were generated using lentiviral transduction similarly as described below, with cell sorting of GFP+ cells to generate stable CAR-expressing cell lines. OKT-3 stock (BioLegend Inc, USA) was diluted to 50mg/ul, then diluted serially threefold. Plates were left at 4°C for 24 hours to bind. Unbound antibody was washed off the plate using PBS. 10^4^ stable Jurkat cells expressing hinge-variant CARs were added to each well; in parallel hinge variant Jurkat-CAR cells were combined with varying doses for SKOV-3 target cells. Plates were incubated at 37°C 5% CO_2_ for 24 hours before staining with mouse anti-human CD69-APC and analysis via flow cytometry.

### Human peripheral blood mononuclear cell (PBMC) isolation

Heparinized whole blood was collected from healthy donors by venipuncture and transported at room temperature from Ottawa Hospital Research Institute. Blood was diluted 1:1 with Hank’s balanced salt solution (HBSS) and PBMCs were isolated by Ficoll-Paque^™^ density gradient centrifugation. Briefly, samples layered on Ficoll-Paque^™^ gradient were centrifuged for 20 min at 700 × g without applying a brake. The PBMC interface was carefully removed by pipetting and was washed twice with HBSS by stepwise centrifugation for 15 min at 300 × g. PBMC were resuspended and counted by mixed 1:1 with Cellometer ViaStain^™^ acridine orange/propidium iodide (AOPI) staining solution and counted using a Nexcelom Cellometer Auto 2000 (Nexcelom BioScience, Lawrence, Massachusetts, USA). T cells from were then activated with Miltenyi MACS GMP T cell TransAct^™^ CD3/CD28 beads and seeded 1×10^6 T cells/ml in serum-free StemCell Immunocult^™^-XF media (StemCell Technologies, Vancouver, Canada) with clinical grade 20U/ml human IL-2 (Novartis)

### Human primary T transduction by spinfection

High concentration lentiviral particles encoding various sdCAR constructs were generated as previously described [29]. After 24 h of T cell stimulation with beads, T cells were transduced with sdCAR-GFP lentiviral vectors (multiplicity of infection = 10) by spinfection. Briefly, lentivirus was added to T cells (1×10^6^ cells/ml) and the mixture was centrifuged at 850 × g for 2 h at 32°C. After centrifugation, cells were incubated at 37°C for another 2 h. After incubation, cells were plated in a 24 well plate (100,000 cells/ml/well in a total of 1.5mL) in StemCell Immunocult^™^-XF supplemented with 20U/mL IL2-2. Media with IL-2 was added at 48 and 72 h post transduction to promote CAR-T cell proliferation without disrupting the cells. Cell number and viability were assessed by AOPI staining and counting using a Nexcelom Cellometer. CAR-T cells were propagated until harvest on days 7, 9, 14, and 21 to assess the efficiency of transduction and to characterize T cell subpopulations by flow cytometry. CAR-T cells that had returned to a resting state (as determined by decreased growth kinetics, day 10 post-T cell activation) were used for assays.

### Continuous live-cell imaging cytotoxicity assay

Cytotoxicity of the CAR-T cells was assayed using a Sartorius IncuCyte^®^ S3 (Essen Bioscience). Tumour cells, U87-MG-vIII-NucLight, MCF7-NucLight, and SKOV3-NucLight or HDF-NucLight, Raji-NucLight, and H292-Nuclight were resuspended in StemCell ImmunoCult^™^-XF with 20 U/mL IL-2 and plated in a flat bottom 96-well plate (2000 cells/well). CAR-T cells or control T cells were added into each well in a final volume of 200 µl per well in StemCell ImmunoCult^™^-XF with 20 U/mL IL-2 at varying effector:target ratios and co-cultured for 7 days at 37°C. Images were taken every 30 min in light phase and under red (ex. 565-605 nm; em. 625-705 nm) or green fluorescence (ex. 440-480 nm; em. 504-544 nm). The assays were repeated thrice with T cells derived from independent blood donors. For one donor, CAR-T cells challenged once or twice with EGFR-high SKOV3 cells were rechallenged with various freshly plated target cells after 7 day of co-culture. Automated cell counting of red (target) or green (CAR-T) cells was performed using IncuCyte^®^ analysis software and data were graphed using GraphPad Prism.

### Animal studies

NOD/SCID/IL2Ry^-/-^ (NSG, JAX #005557) mice were purchased from Jackson Laboratories and maintained by the Animal Resource Group at the National Research Council of Canada.. Eight-week-old NSG mice were injected with 2×10^6^ SKOV3-NucLight in 100 ul of Hanks’ Balanced Salt Solution (HBSS) subcutaneously. Eighteen days post tumour injection (when tumour reached approximately 5mm × 5mm), mice were retro-orbitally injected with 1×10^7^ fresh day 10 post activation mock T cells or T cells transduced with various CAR-T cells as described in the text. Tumours were measured using calipers twice a week and mice were imaged via IVIS *in vivo* imager for red-fluorescence signal (expressed on tumour cells) once a week. For the alternative U87 MG vIII model experiments, mice were subcutaneously injected with 1×10^6^ fluorescently labelled U87 MG-vIII cells described above, a number we previously determined to consistently produce a palpable tumour within 7 days. Eight days after tumour cell injection, cryo-preserved CAR-T cells were thawed, washed with PBS, and 5×10 total T cells (with 20-25% CAR transduction) were immediately delivered intratumourally, ensuring equal distribution of tumour sizes between groups. Tumour growth was evaluated three times per week using calipers by trained animal technicians blinded to specific treatment groups. Mice were also assessed for tumour growth using IVIS *in vivo* imaging to examine red fluorescence derived from the NLS-mKate2 marked U87-MG-vIII cells. In order to minimize requirement for animal shaving to the area immediately around the implanted tumour a blanket was used to obscure part of animals during imaging. Primary endpoint was tumour size above 2000mm^3^, with secondary endpoints determined by overall animal health and well-being. Mice were euthanized when they met pre-specified endpoints. The study was approved by the NRC-HHT Institutional Animal Care Committee and was conducted in accordance with Canadian Council on Animal Care (CCAC) guidelines. Tumour growth and survival (humane endpoint) curves were generated using GraphPad Prism.

### Generation of a monoclonal anti-sdAb antibody for assessment of sdCAR surface expression

CAR expressing Jurkat cells or primary T cells were generated as indicated elsewhere and assessed for surface expression using an Alexa-Fluor647 labelled murine monoclonal anti-sdAb antibody generated in house. In brief, mice were immunized with sdAb fragments produced in e coli followed by B cell isolation and hybridoma formation as previously reported[32]. Clonal hybridoma supernatants were screened for reactivity against a number of plate bound sdAbs to identify broadly sdAb-reactive monoclonal antibodies. Antibodies were then sequences and transferred to a recombinant murine IgG2a backbone in order to ease production in CHO cells. Antibodies were then purified via protein-G column before labelling with Alexa-Fluor 647 using a commercial kit. This reagent was confirmed to react broadly against sdAb-CAR expressing cells but not scFv-CAR expressing Jurkat cells. This reagent was then employed through cell staining and flow cytometric assessment as described in the text for quantification of relative sdCAR surface expression.

### Flow cytometry and antibodies

For in vitro studies, cells were stained antibodies as indicated in the text, incubating the cells for 15 minutes at room temperature before washing with PBS. Cells were then resuspended in PBS and examined using a BD Fortessa Flow cytometer. Staining of human EGFR was performed using anti-human EGFR-PE-CF594 (BD Biosciences, Cat #563431), and anti-sdCAR staining was performed using in house antibody generated as described above. Blood was obtained from mice at various time points post CAR-T injection. Blood was washed with cold phosphate-buffered saline (PBS) and pelleted at 350 × g for 5 min at 4°C. Red blood cells were lysed using Red Blood Cell Lysing Buffer Hybri-Max (Sigma-Aldrich, St. Louis, MO, USA). Human T cells were identified and analyzed for activation/differentiation status using the following antibodies: hCD45-APC-H7, hCD45RA-BV650, hCD45RO-PE-CF594, hCD27-BUV737, hCCR7-PE, hCD4-BUV395, and hCD8-PerCP-Cy5.5 (all antibodies from BD Biosciences, USA). CAR expression was measured indirectly via expression of GFP incorporated in CAR constructs. To evaluate exhaustion, staining by an hPD-1-BV421 antibody was evaluated. T cell activation was detected using hCD25-PE-Cy7 and hCD69-BV786 antibodies. For *in vivo* studies, a BV711-labeled antibody against mouse CD45 was used to identify murine cells.

## Results

### High-affinity EGFR-specific sdAbs generate antigen-responsive CARs

Generation of camelid single domain antibodies (sdAbs) against EGFR using DNA immunization and phage display was reported in a previous study [28] (see Supplementary Fig. 6 for an overview of the workflow employed here). In order to assess whether these sdAbs were functional in the context of a CAR molecule we chose three EGFR-specific sdAbs with varying affinities and epitopes (Table 1)[28]. The sdAbs were cloned into a modular CAR backbone and the resulting EGFR-sdCARs were screened for responses to target cells with varying EGFR expression (supplemental Fig. 1) using a previously described high throughput CAR-Jurkat screening assay[29] (Fig. 1A). Jurkat cells electroporated with any of three EGFR-specific sdCAR constructs showed specific upregulation of CD69 following co-culture with EGFR-high SKOV3 cells or EGFR-low MCF7 cells (Fig. 1B,C), but no activation in response to EGFR-negative Raji cells (Fig. 1D). To investigate whether these EGFR-sdCAR constructs might also induce on-target off-tumour responses against non-malignant cells, we tested a similar CAR-J assay against human dermal fibroblast cells (HDF) from a healthy donor, which showed high levels of EGFR expression (Supplemental Fig 1). In this case, we observed strong anti-HDF responses for Jurkat cells transiently expressing any of the EGFR-sdCAR constructs tested (Figure 1E). These data demonstrate that high affinity EGFR sdAb-based CAR molecules can induce strong responses to EGFR-expressing target cells.

**Figure 1:**
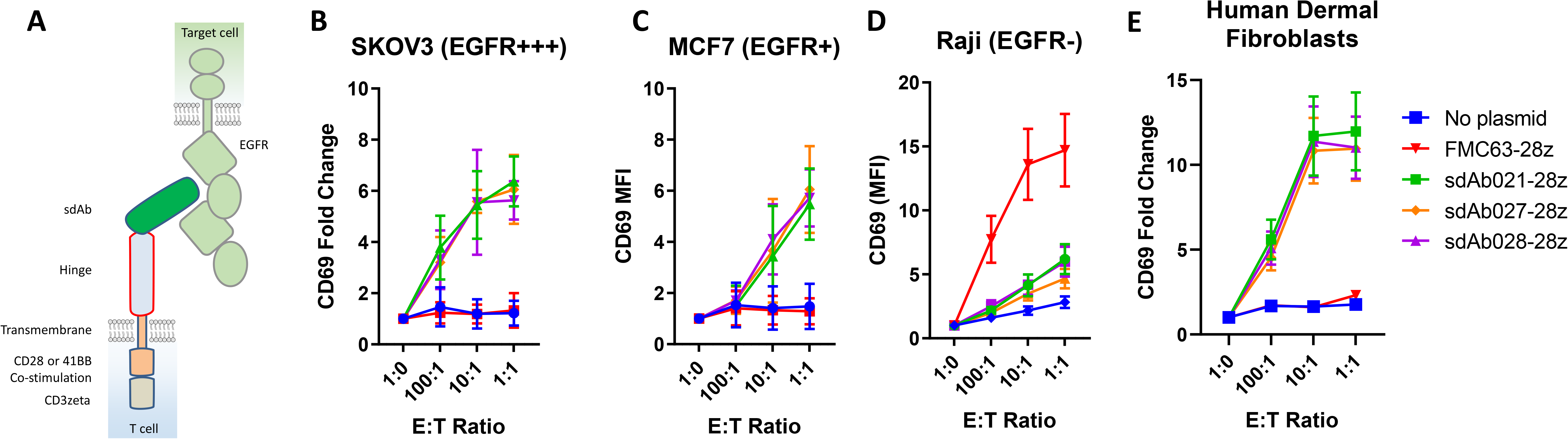
Identification of EGFR-specific sdAbs with CAR functionality. (A) The structural elements of human EGFR and sdAb based EGFR-targeting CAR tested here are shown. Various EGFR-specific sdAbs were cloned into a modular CAR backbone via golden gate cloning. Jurkat cells were then electroporated with the resulting constructs. Control cells with no plasmid or with a CD19-targeted scFv element were also tested here (FMC63-28z). Jurkat cells (30 000/well) transiently expressing various CAR plasmids as shown were co-cultured with varying doses of (B) EGFR-high SKOV3 cells, (C) EGFR-low MCF7 cells, (D) EGFR-negative Raji cells, or (E) EGFR-high healthy donor derived human dermal fibroblast cells, and examined for activation via staining with APC-labelled anti-human CD69 antibody. Results show the mean +/- SEM from a three independent experiments performed in duplicate.

**Table 1.**
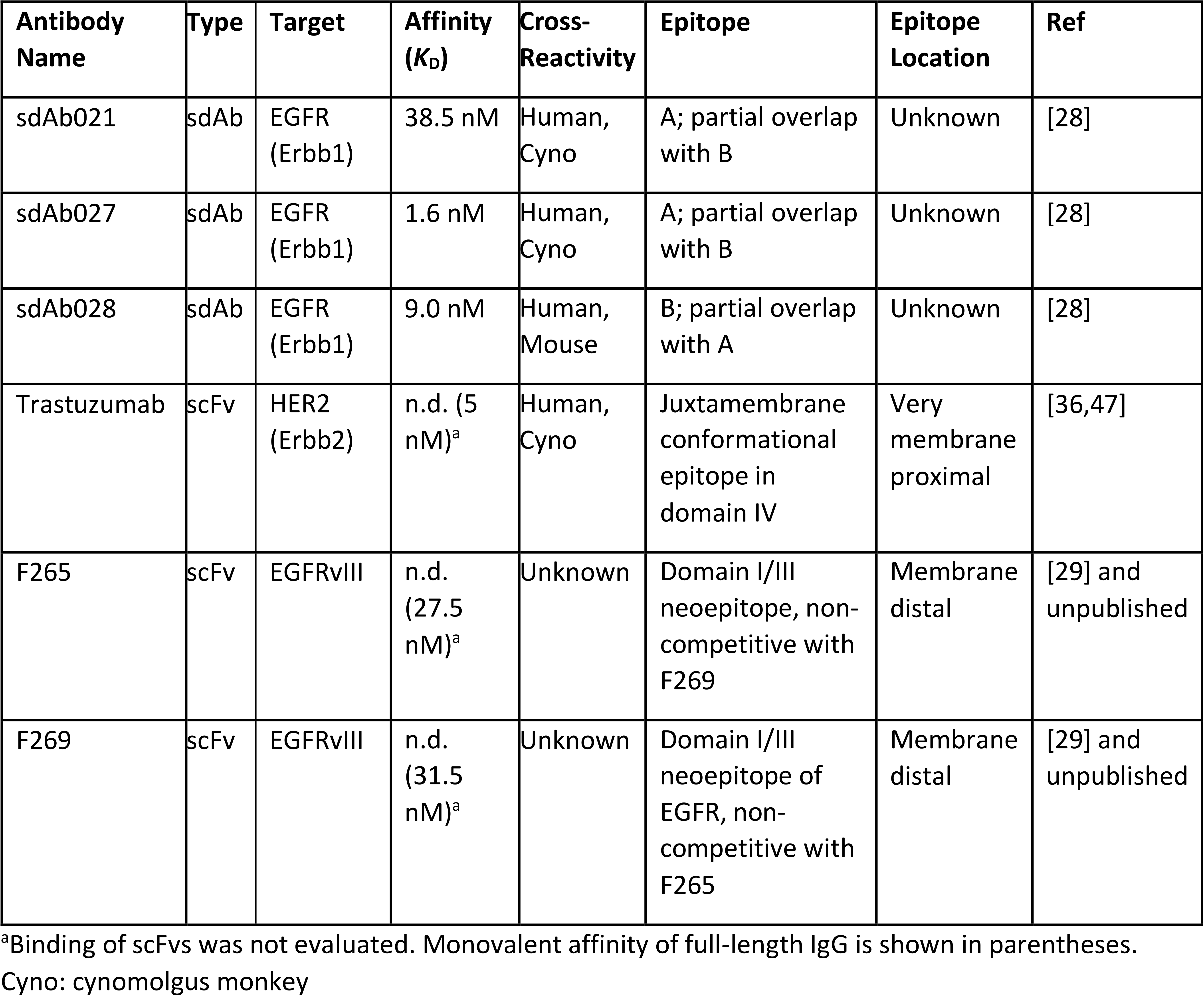
Antigen-binding domains tested as CAR elements in this study.

### Hinge truncation decreases CAR antigenic sensitivity

It has previously been shown that the size of the spacer/hinge domains used in a CAR can have a strong effect on CAR activity[33,34]. We hypothesized that rather than using variably sized hinge domains derived from various human proteins, it may be possible to simply truncate the hinge region to adjust target sensitivity. We cloned EGFR sdCAR constructs with either a full-length 45 amino acid human CD8 hinge (45CD8h) or N-terminally truncated hinge variants (34CD8h, 22CD8h, or no hinge; see Fig. 2A). Jurkat cells transiently expressing the truncated hinge variants of all three EGFR sdCARs showed progressively decreasing activation in response to SKOV3 target cells with high EGFR expression (Fig. 2B-D). One possible explanation of the consistent effect of hinge truncation on antigenic sensitivity might be a reduction in surface expression for hinge-truncated CARs. Using a broadly reactive anti-sdAb antibody to label surface sdCAR expression we observed no correlation between hinge domain and CAR surface expression (Supplemental Fig 2A,B). We also confirm that hinge-truncated CARs are not somehow inhibiting global Jurkat responsiveness as hinge variant sdAb-CARs show similar response to plate-bound anti-CD3 (Supplemental Figure 2C, D).

**Figure 2:**
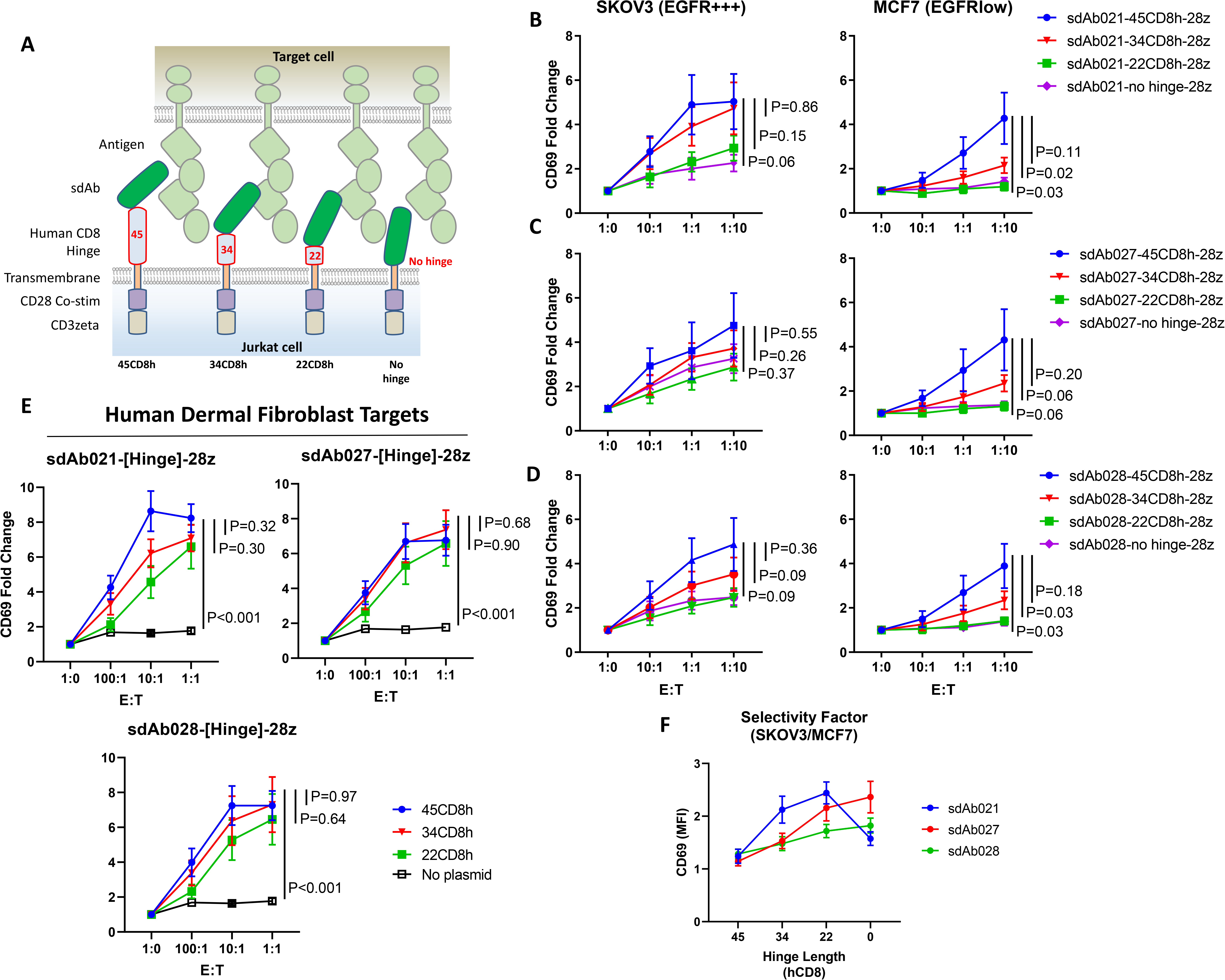
Hinge truncation decreases target response for EGFR-sdAb CAR constructs. (A) Single domain antibody based CARs targeting human EGFR were generated with hinge domains of varying length [full length human CD8-hinge (45CD8h), truncated CD8-hinge (34CD8h or 22CD8h), or no hinge element. (B-D) Jurkat cells were electroporated with varying CAR constructs before co-incubation with no target cells (1:0 E:T), EGFR-high SKOV3 cells, or EGFR-low MCF7 cells, (E) similarly hinge-variant EGFR-CAR-Jurkat cells were co-incubated with human dermal fibroblast cells. After overnight incubation CAR/GFP+ cells were examined for activation via staining with APC-labelled anti-human CD69 antibody. (F) Shows the relative CAR-J activation of various constructs in response to SKOV3 or MCF7. Results show the mean +/- SEM from a single experiment performed in duplicate. P values are derived from a student T test comparing 45CD8h and other CARs tested in parallel at 1:1 E:T

Next, we tested the response of EGFR sdCARs with truncated hinge domains against EGFR-low MCF7 cells, where we observed similarly decreased response for hinge-truncated EGFR-sdCARs. Interestingly, response to EGFR-low MCF7 cells appeared more sensitive to hinge truncation than for EGFR-high SKOV3 cells, particularly for the sdAb021 sdCAR (Fig. E, Supplementary Fig. 2C). We next performed high-throughput screening of hinge-truncated CARs against EGFR-high fibroblast cells derived from healthy human donors, where we observed high CAR activity with progressively decreasing response for truncated hinge constructs (Fig 2E). Comparing the relative response to EGFR-high SKOV3 over EGFR-low MCF7 cells we note that the highest selectivity with sdAb021-based EGFR-sdCARs and thus selected this construct for further testing described below (Fig. 2F). Overall, these results indicate that hinge truncation can be an effective tool to adjust antigenic-sensitivity of EGFR-sdCARs.

### Single Amino Acid Hinge Truncation Provides Fine-Tuned Control of CAR-Antigen Response

Given our initial observations with truncated forms of the CD8-hinge domain, we wanted to more finely map the effects of hinge truncation on an EGFR sdAb CAR. Given the effects of hinge truncation, we reasoned that hinge lengthening might be beneficial for antigen response; thus, we designed an extended hinge domain wherein an additional N-terminal flexible linker of 17 amino acids were appended to the human CD8-hinge sequence [(GGGGS)_3_GG-45CD8h] within the EGFR-sdCAR construct. We then generated a library of sdAb021-CAR constructs with single amino acid N-terminal deletions of the lengthened human CD8 hinge (every combination between 62 and 1 amino acid). Screening our sdCAR single-residue hinge truncation library revealed a clear pattern of CAR activation (Fig 3A, Supplemental Fig 3A). CAR constructs containing a full human 45AA CD8-hinge or longer produced a significantly higher response to SKOV3 cells and a lower but consistent response to EGFR-low MCF7 cells. While there was some variation, the addition of a flexible linker extension to the CD8-hinge domain did not seem to have a positive or negative effect on CAR response. EGFR-sdCARs with CD8 hinge sizes between 45 and 26 amino acids showed a progressive decrease in CAR activation, while CAR constructs with hinges 26 or less amino acids within their hinge domains showed no response to EGFR-expressing targets over CAR-J cells in the absence of targets. As in experiments above, partial truncation of the hinge domain (less than 40 amino acids) resulted in an EGFR-sdCARs with sustained response to EGFR-high SKOV3 cells but similar lack of response to EGFR-low MCF7 or EGFR-negative Nalm6 target cells. Taken together, these data suggest that hinge truncation can be used for precise control over EGFR-sdCAR antigenic sensitivity.

**Figure 3:**
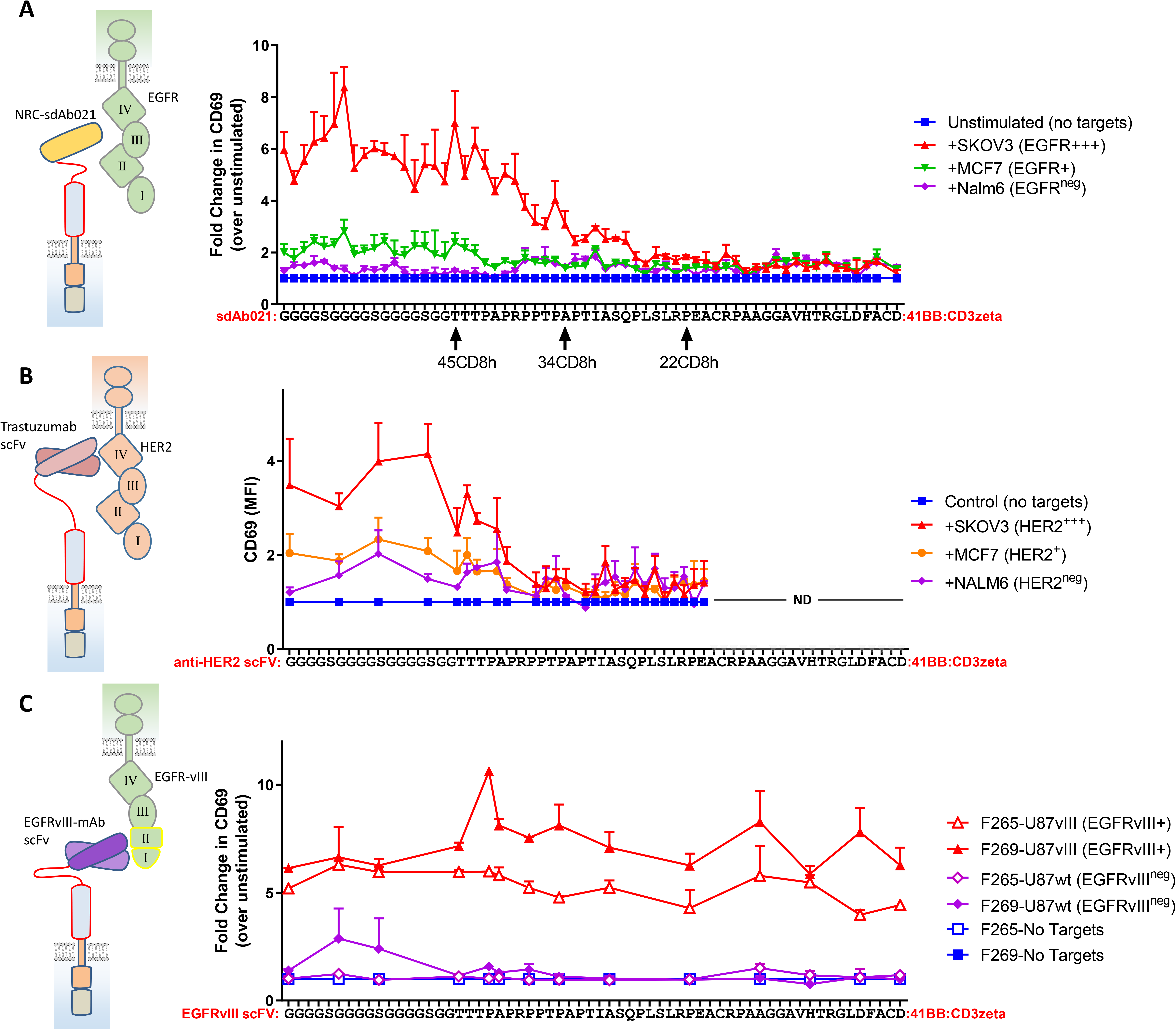
CAR hinge sensitivity is highly dependent on target antigen and epitope location. (A) A range of sdCAR constructs were produced containing the EGFR-specific sdAb021 and human CD8 hinge domains ranging between 1 and 45 residues in length. The full-length 45-residue CD8 hinge was extended with a Gly-Ser linker between 1 and 17 residues in length. Jurkat cells were electroporated with the resulting constructs and then co-incubated at an effector to target ratio of 1:1 with EGFR-high SKOV3 cells, EGFR-low MCF7 cells, or EGFR-negative NALM6 cells. After overnight incubation, CAR/GFP+ cells were examined for activation via staining with APC-labelled anti-human CD69 antibody. (B) Similarly, trastuzumab-scFv HER2-specific CAR constructs were generated with hinge domains of varying lengths. CAR-expressing Jurkat cells were co-incubated with HER2-high SKOV3 cells, HER2-low MCF7 cells, or HER2-negative NALM6 cells and then CAR-T activation was assessed. (C) Similarly, EGFRvIII-specific CAR constructs were generated with hinge domains of varying lengths. CAR-expressing Jurkat cells were co-incubated with EGFRvIII-overexpressing U87vIII cells or EGFRvIII-negative U87wt cells and then CAR-T activation was assessed. Results show means +/- SEMs of three separate experiments.

### Epitope location is critical determinant of CAR hinge sensitivity

As has been previously proposed, the location of the CAR target epitope should be critical in determining the minimal hinge size needed for CAR antigenic responsiveness[35]. We do not have definitive data as to the epitope(s) targeted by the EGFR sdAbs used in our sdCAR constructs, though epitope binning experiments indicate that all three sdAbs have partially overlapping epitopes, and cross-reactivity analysis is suggestive of membrane proximal domain IV binding[28] (Table 1). To more carefully investigate the role of epitope location, we generated limited hinge libraries for CARs with known target epitopes. Using scFv-CARs based on either trastuzumab, which is known to bind a highly membrane proximal epitope of HER2 [36], or novel EGFRvIII-specific antibodies[29], which by necessity must bind the membrane distal neo-epitope of EGFRvIII (Table 1). The membrane-proximally targeted trastuzumab-derivative scFv-CAR required a very long hinge element; no HER2 scFv CARs containing shorter than a full CD8-hinge responded against HER2+ target cells (Fig. 3B, Supplemental Fig 3B). In contrast, membrane-distal targeting EGFRvIII CARs maintained full activation with all hinge formats, even when the entire CD8 hinge domain was deleted (Fig. 3C, Supplemental Fig 3C). These data support previously proposed models wherein epitope location is a key determinant of the hinge requirement for CAR functionality, and thus hinge truncation can only be employed for fine-tuning antigenic sensitivity in membrane-proximal targeting CAR constructs.

### Hinge-truncated CAR responses is consistent in primary T cells *in vitro* and *in vivo*

We next undertook extensive experiments to confirm whether the reduced response of sdCARs with truncated hinge elements observed in Jurkat cells was similar in the responses of primary CAR-T cells. Thus, T cells were isolated from whole blood obtained from three healthy human donors and transduced with CAR lentivirus encoding three hinge-variant forms of sdAb021-41BB-CD3z sdCAR. Following polyclonal expansion and sdCAR transduction, green fluorescent protein (GFP+) CAR-T cells were placed in low density co-culture with target cells stably marked with a nuclear localized red fluorescent protein. As observed in Jurkat cells, CAR-T surface labelling using an anti-single domain antibody did not reveal any correlation between hinge length and CAR surface expression (Supplemental Figure 4). Co-cultures were then monitored for tumour cell (red fluorescence) and CAR-T cell (green fluorescence) growth over 7 days without media changes. Consistent with observations in Jurkat cells, hinge truncation progressively diminished the ability of primary sdAb021 sdCAR-T cells to restrict EGFR-high SKOV3 tumour cell growth and induce CAR-T cell expansion (Fig. 4A,B, Supplemental Fig 5), although all hinge formats maintained increased SKOV3 killing relative to untransduced cells (mock). We next investigated how hinge-truncated EGFR-sdCARs would respond to EGFR-low target cells. Again consistent with Jurkat observations, only EGFR-sdCAR-T cells with the unmodified CD8-hinge domain (sdAb021-45CD8h-28z) showed potent target killing and CAR-T expansion in response to EGFR-low MCF7 cells (Fig. 4C,D, Supplemental Fig. 5).

**Figure 4:**
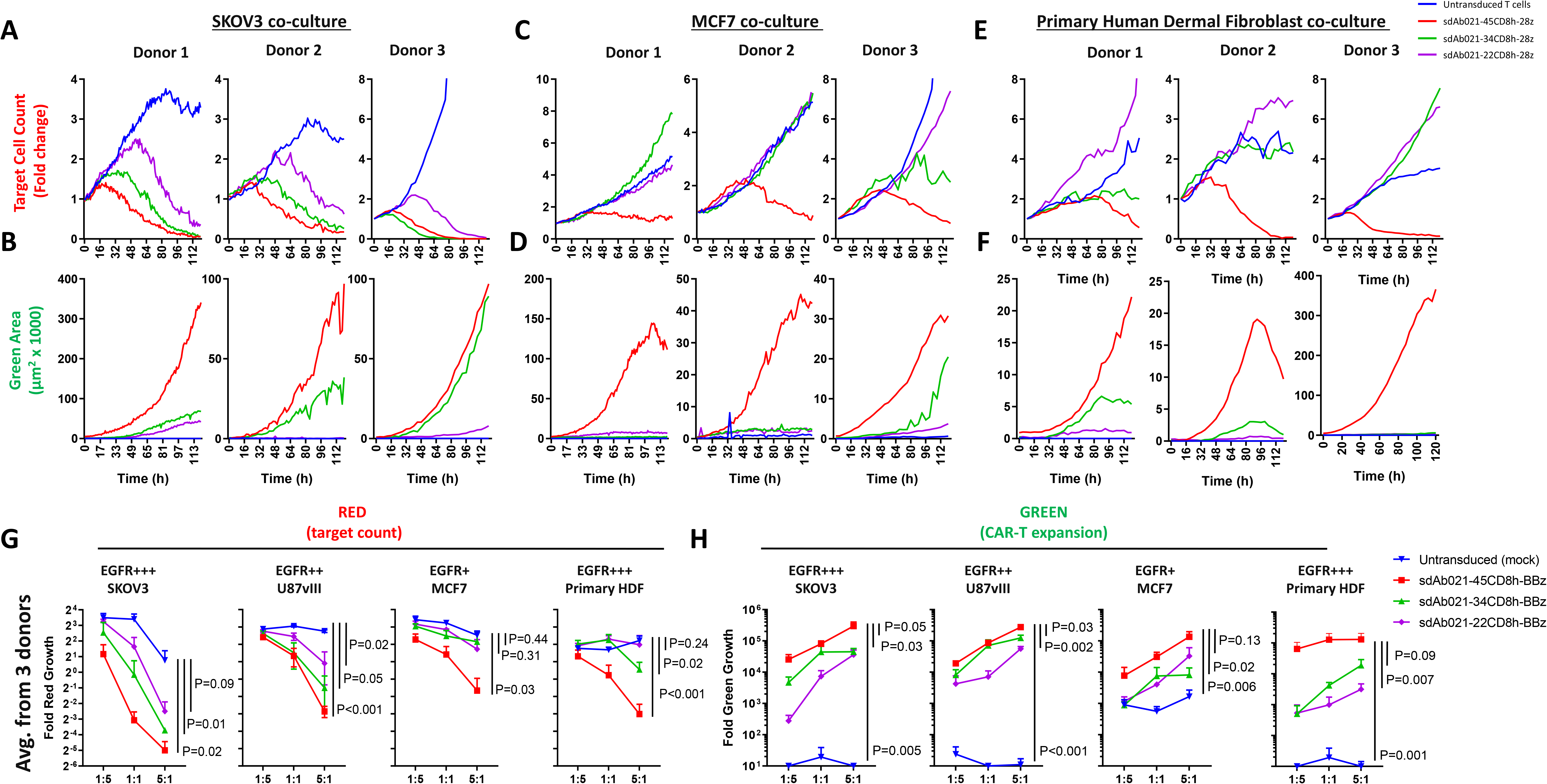
Hinge truncation progressively diminishes tumor cell killing and expansion of primary sdCAR-T cells in response to EGFR expressing cells. Concentrated lentiviral particles encoding hinge-modified EGFR-specific sdAb021 CARs as well as a GFP marker were generated. Peripheral blood T cells were isolated from two donors before polyclonal expansion and lentiviral transduction. Varying doses of sdCAR-T cells or mock transduced cells (empty CAR backbone lentivirus) were added to a 96-well plate containing 2000 EGFR-high SKOV3, EGFR-medium U87vIII, or EGFR-low MCF7 target cells stably expressing nuclear localized mKate2 (red). Co-cultures were examined over 7 days via live fluorescence microscopy (Incucyte) to differentiate red-fluorescent target cell counts or total area of green-fluorescent CAR-T cells. (A-B) Depicts the response to EGFR-high SKOV3 targets, (C-D) depicts the response to EGFR-low MCF7 targets, (E-F) depicts the response to EGFR-high human dermal fibroblast cells. Each graph depicts automated cell counts or fluorescent areas from a single well from a single experiment. (G) Day 5 mean fold change in target cell growth at varying E:T ratio for hinge-variant EGFR-sdCAR-T cells across 3 donors is shown +/- SEM, (H) similarly the mean fold expansion of hinge-variant EGFR-sdCAR-T cells from 3 donors is shown +/-SEM. P values are derived from a student T test comparison on 45CD8h-hinge containing constructs with other constructs at an E:T of 5:1.

In order to investigate the behaviour of EGFR-sdCAR-T cells in the context of non-malignant EGFR-expressing cells, we performed a similar co-culture assay using human dermal fibroblast (HDF) cells derived from a healthy human donor. As observed in SKOV3 and MCF7 cells, hinge truncation results in progressively decreased target killing and CAR-T expansion in response to HDF targets (Fig. 4C,D). Plating EGFR-hinge variant CAR-T cells with target cells at varying effector:target ratios revealed a consistent pattern of enhanced selectivity for EGFR-high targets (SKOV3 or U87vIII) over EGFR-low tumour targets (MCF7) or healthy-donor HDF cells [Figure 4E,F]. Interestingly, hinge truncated EGFR-sdCAR constructs seems to increase selectivity for EGFR-high SKOV3 or U87vIII tumour cells over HDF cells, despite relatively similar EGFR expression (Supplemental Fig 1); underlining the complex nature of CAR-T target cell recognition. Looking at the compiled data from 3 donor CAR-T cell products, we observe that only the full length CD8-hinge containing EGFR-sdCAR-T cells induce significantly more tumour killing than mock T cells (Figure 4G). Interestingly, expansion of EGFR-sdCAR-T cells with truncated hinge domains seems to be somewhat less affected by hinge truncation, although all constructs are significantly diminished in response relative to full length hinge (Figure 4H). Overall these results provide strong evidence that that hinge truncation can be used to increase CAR selectivity for EGFR-overexpressing tumours, and decrease potential on-target off-tumour effects.

We also investigated the effect of hinge truncation on serial CAR-T killing. We isolated hinge modified sdCAR-T cells following primary challenge with antigen-overexpressing SKOV3 cells using the low density co-culture assay as described above and re-challenged the sdCAR-T cells by re-plating with additional target cells (Supplemental Figure 6). Re-challenged cells maintained their relative selectivity observed in primary challenge as determined by target killing and CAR-T expansion, which both decreased with hinge length. These results indicate that re-stimulated CAR-T cells show similar or higher antigenic discrimination as observed in primary challenge, with progressively decreasing antigen-induced expansion following serial challenge.

### *In vivo* CAR-T response is progressively diminished with hinge truncation

Finally, we wished to investigate whether our *in vitro* observations regarding the relationship between hinge length and CAR-T activity were consistent in an *in vivo* xenograft model. Using the relatively slow growing SKOV3 model, mice were challenged with 2×10^6^ SKOV3 tumour cells subcutaneously. After allowing 18 days for tumour implantation, mice with palpable tumours were injected with 10 million hinge modified sdCAR-T cells or mock transduced CAR-T cells. There was a progressive increase in tumour growth in all mice (Fig. 5A) and decreased survival was observed in mice treated with hinge-truncated sdCAR-T cells (Fig. 5B). Surprisingly, mice treated with sdCAR-T cells bearing the shortest hinge (sdAb021-22CD8h-28z) showed apparent decreased survival compared with untreated animals, although larger experiments would be needed to confirm this result. The progressive effect of hinge truncation could be clearly observed in the final tumour volume measurement taken at day 112 post tumour challenge (Fig. 5C,D) prior to early experimental termination due to non-experimental facility disruptions. Only those mice treated with the longest hinge format sdAb021-45CD8h-BBz showed a significant decrease in tumour volume relative to untreated mice, but a progressive decrease in CAR activity with CAR-T cells expressing truncated hinge is nonetheless readily apparent (Figure 5C).

**Figure 5:**
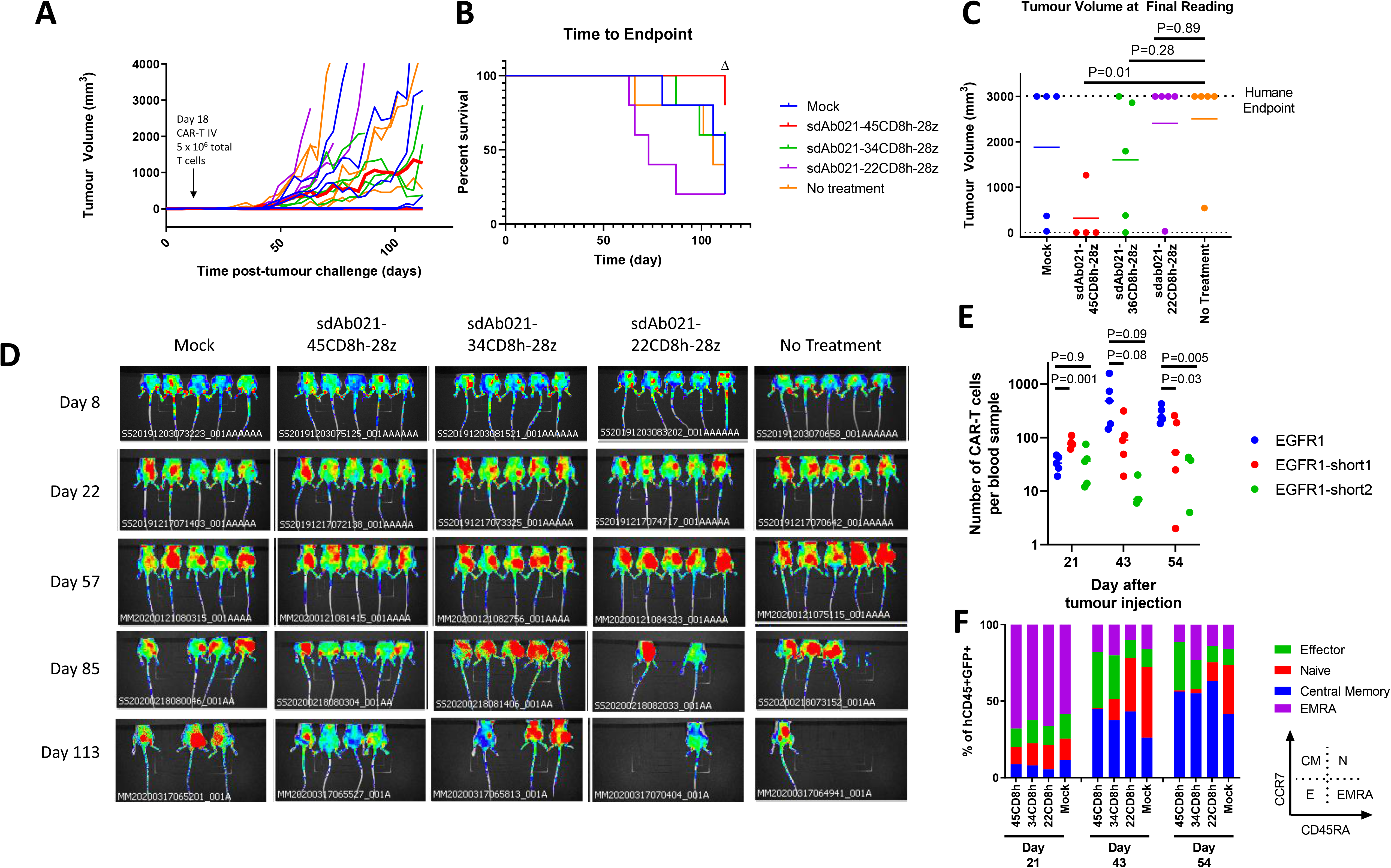
Hinge truncated EGFR sdCAR-T cells show progressively diminished response to target cells *in vivo*. NOD/SCID/IL2γ-chain-null (NSG) mice were injected subcutaneously with 2×10^6^ SKOV3 cells stably expressing mKate2. Mice were injected with 5×10^6^ total T cells (∼1×10^6^ CAR-T cells) intravenously. (A) Tumor volume was estimated using caliper measurements and (B) time to defined humane endpoint (tumor volume 2000 mm^3^) was assessed. Δ Note that the experiment was ended early due to non-experimental related animal facility shutdown, and thus the final tumor measurement of day 112 is shown in (C). (D) Fluorescence imaging was performed at varying timepoints after tumor cell injection. (E) Blood was also collected at selected timepoints after tumor challenge to quantify the proportion of human CD45+ cells that were GFP/CAR+. (F) Staining for human CCR7 and CD45RA was used to assess the differentiation status of CAR/GFP+ cells or total T cells for mice treated with mock-transduced CAR-T cells. See inset for gating strategy used to delineate effector, naïve, central memory, or effector memory-RA+ (EMRA) cells.

Examining CAR-T cells in the blood of treated mice revealed a consistent pattern of increased expansion of sdCAR-T cells with longer hinge regions at 43 and 54 days post-tumour injection (Fig. 5E). Analysis of T cell differentiation revealed increased naïve sdCAR-T cell and decreased effector sdCAR-T cell populations were for hinge truncated sdCARs (Fig. 5F), supporting the interpretation that truncated sdCAR-T cells show progressively decreased in *in vivo* responses. We also performed a similar experiment in which wild-type or hinge modified sdCAR-T cells were delivered intratumourally in the relatively more aggressive U87-vIII glioblastoma xenograft model. We observed similar progressive decreases in anti-tumoural effect (Fig. 6A-C) and CAR-T expansion (Fig. 6D) and progressive increases in naïve CAR-T cell populations associated with hinge truncation (Fig. 6E). Taken together, these results demonstrate that hinge truncation can be used to precisely program antigenic sensitivity of CAR-T *in vitro* and *in vivo*.

**Figure 6:**
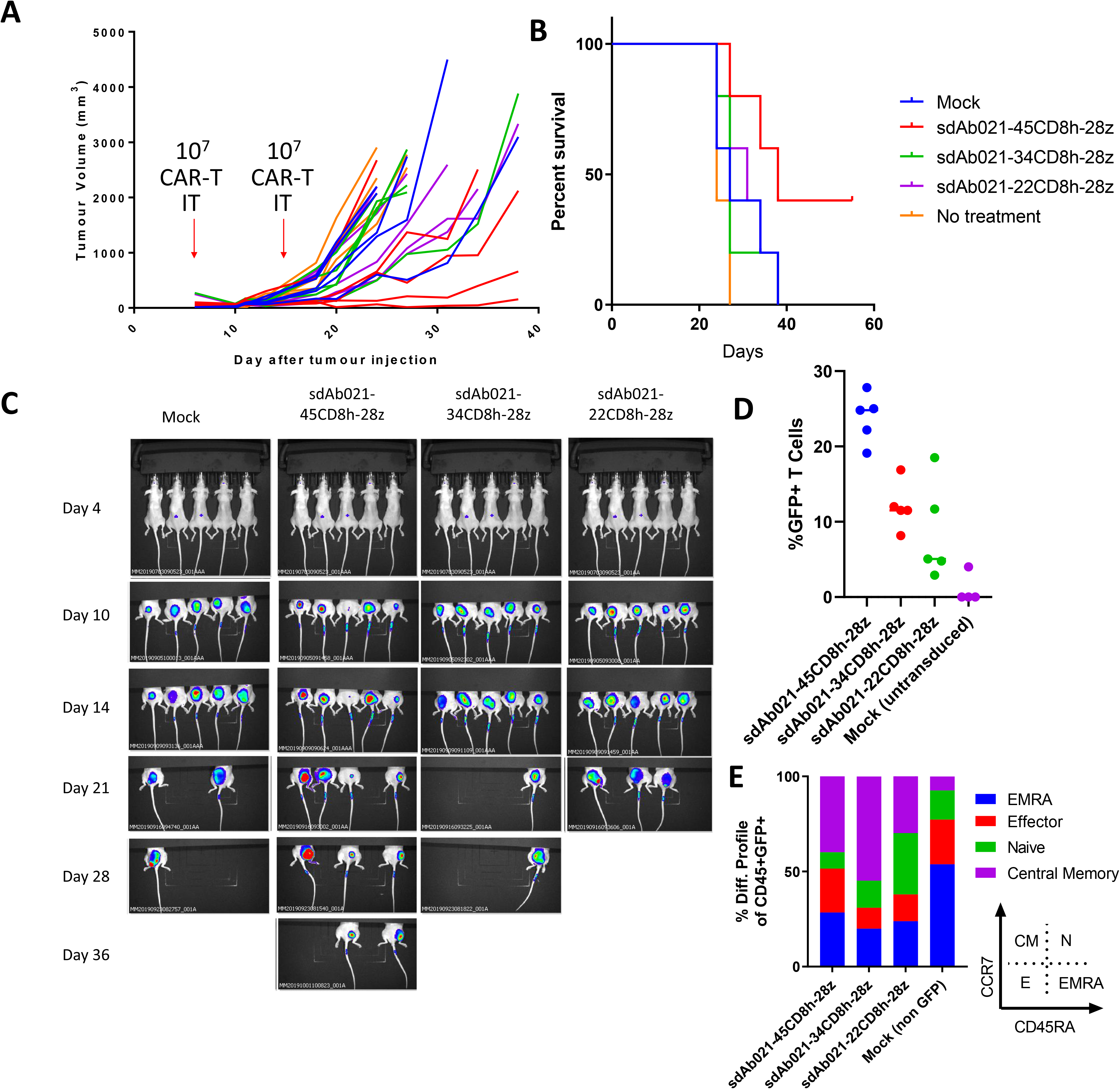
Hinge truncation results in similar progressively diminished EGFR-specific sdCAR activity in the more aggressive U87vIII xenograft tumor model. (A) NSG mice were injected subcutaneously with 1×10^6^ U87vIII cells stably expressing mKate2. At 7 and 14 days post-tumor challenge mice were injected intratumorally with 1×10^7^ total T cells (approx. 2.5×10^6^ hinge-modified sdCAR-T cells) or with untransduced control T cells (mock). Tumor growth was monitored via caliper measurements. N=5 mice per group. (B) Mice were sacrificed at pre-determined endpoints based on animal condition or tumor volume >2000 mm^3^. (C) *In vivo* imaging was performed to examine the mKate2 fluorescent signal associated with tumor cells. (D) Blood was drawn from challenged mice and examined for the proportion of sdCAR-transduced cells (GFP+) within the hCD45+ lymphocyte fraction at the final timepoint where all experimental mice were alive (day 21). (E) sdCAR-T cells, or total hCD45+ cells for mock T cell treated mice, were examined for differentiation status using antibody staining for human CCCR7 and CD45RA. See inset for gating strategy used to delineate effector, naïve, central memory, or effector memory-RA+ (EMRA) cells.

## Discussion

We sought here to develop a novel EGFR-specific CAR construct that can discriminate between high level expression on tumours and lower expression on normal cells. Despite some variation in binding affinity for the three EGFR sdAb moieties tested here (ranging from 1.6 to 38 nM; see Table 1), all three sdCAR constructs showed similar strong responses to both EGFR-high SKOV3 and EGFR-low MCF7 target cells. Interestingly, we note EGFR-sdCAR responsiveness to MCF7 cells despite no apparent reactivity of the purified sdAbs to EGFR-low MCF7 cells previously reported [28]. These results are consistent with previous observations that EGFR specific CARs are relatively insensitive to ABD affinity up to the micromolar range[27] and underscore the exquisite antigen sensitivity of CAR-T cells to respond to and lyse even very low antigen expressing target cells[37–39]. This phenomenon possibly relates to the extreme multi-valency of both CAR and EGFR on their respective cells; increased valency is well known to lead to significant avidity effects that boost the apparent affinity of biological interactions[30,40–42], and these would presumably apply to CAR-T cells as well. These results point to a possibility that perhaps lowering ABD affinity might be a somewhat blunt tool for modulation of CAR antigenic sensitivity.

Previously, affinity modulation has been used to increase selectivity of CAR constructs for overexpressing cells[27]. While it is certainly possible to decrease sdAb affinity through various molecular strategies[43], mutational antibody changes can sometimes lead to unpredictable binding behaviour such as unexpected off-target binding, elevated tonic signaling, or poor protein stability. Thus, we wished to pursue an alternate strategy to decrease on target activity through hinge modification. The use of hinge domains derived from various antibody isotypes or receptor ectodomains has been well- documented to have powerful influence on antigenic responsiveness of particular CAR constructs[2], and the strategy to employ hinge domains of varying length has also been previously explored[3,33,44,45]; to our knowledge however, this is the first report of a simple truncation strategy of the human CD8-hinge domain which is most commonly employed in CAR designs.

We find that truncations of a human CD8 hinge by as little as a single amino acid can be a remarkably precise tuning mechanism for CAR antigenic sensitivity. For the EGFR sdCAR we tested most extensively here, there was a steady drop-off of antigenic CAR response over a range of 10 to 20 amino acids within the CD8 hinge motif. While the requirements of lentiviral production and primary T cell transduction dictated that we could only test a limited number of constructs in primary T cells, it would be intriguing to map the optimal hinge length for antigen induced CAR-T killing or CAR-T expansion in genuine donor-derived T cells. Our hinge-truncation data using either membrane proximal (trastuzumab) or membrane-distal (anti-EGFRvIII-mAbs) based CAR constructs provides a further molecular demonstration of the criticality of epitope location as a determinant of hinge-sensitivity for CAR molecules reacting to tumour cells.

While the exact epitope for the EGFR sdAbs tested is not known, the data here would indicate a likely membrane proximal location. Consistent with this, sdAb028 cross-reacts with human and mouse but not cynomolgus EGFR; the only positions at which mouse and human EGFR sequences are identical but diverge from cynomolgus EGFR are in domain IV of EGFR[28]. Our finding that a full length CD8 hinge is required for the trastuzumab scFv-CAR seems to be consistent with previous experiments using similar CARs where hinge domains were also included[46].

In contrast to hinge truncation, we found that extension of the CD8 hinge domain does not have a strongly positive or negative effect on CAR response, at least as determined by the CAR-J assay. Similarly, membrane-distal epitope targeting EGFRvIII CARs showed similar responses across all hinge formats tested here. Previous work has indicated that longer hinges can decrease *in vivo* activity for membrane-distal epitope targeting CARs[44], but follow-up studies pinpointed the effect to be related to FC-binding by IgG-hinge motifs rather than hinge length specifically[45]. It is also important to note that the multi-[GGGS] linker format used to extend our hinge is a highly flexible sequence typically employed for scFv engineering, and thus may allow for adequate ABD motility and close CAR packing unlike long hinge domains used in previous studies [33,44]. While certainly membrane-distal CAR molecules cannot functional with a hinge domain which is too short, no firm conclusion on the impact of an excessively long hinge can yet be drawn.

Due to the relatively more demanding technical requirements of testing CAR-T constructs in primary T cells we only tested a limited number of CAR constructs within primary cell assays. Nonetheless, data presented here provide additional evidence that molecular optimization using transient CAR expression in Jurkat cells is predictive of responses in stably transduced primary CAR-T cells, as we have previously reported[29]. The wider use of such optimization methodology could lead to improved ABD/hinge design for future CAR products. It may be possible for instance to design CARs with customized signaling for application in CD4, CD8, gamma-delta T cells, or NK cells.

The *in vivo* experiments presented here also underline the fundamental trade-off between on-target, on-tumour activity and strategies that to reduce on-target off-tumour responses. Achieving a level of CAR signaling that is adequate for potent tumour killing yet also eliminates the risk of on-target toxicity may be a difficult balance. For mice treated with the shortest hinge format tested, we observed a trend towards shorter time to humane endpoint, perhaps reflecting CAR-T expansion without tumour cell killing, something that is also apparent *in vitro* CAR-T experiments. While the partially hinge truncated EGFR-sdCARs that showed the best SKOV3-selectivity *in vitro* (sdAb021-34CD8h-BBz) had relatively little therapeutic effect *in vivo* here, it may be possible to combine such reduced sensitivity hinge-truncated EGFR-sdCAR in a multi-antigen targeting CAR strategy that will ultimately recognize and lyse tumour cells in a more selective fashion. Importantly, the aggressive xenograft models used here are imperfect and would likely not perfectly predict functional CAR-T responses in humans with slower growing and/or metastatic disease.

While CAR constructs with maximal antigen sensitivity might be desirable in certain contexts, such as for CAR-T therapies targeting B cell family restricted antigens, the presence of EGFR expression on normal tissue requires a targeting strategy that would be selective for EGFR-overexpressing cancer cells. The data presented here provide strong evidence that hinge truncation can be a powerful tool to engineer CAR molecules with desired antigenic sensitivity, for potential downstream development in multi-antigen targeting CAR therapeutics. The clinical use of CARs with varying hinge lengths could also present an intriguing alternative safety pathway for clinical trials of solid tumour targeting CARs through hinge length escalation.

## Declarations

- The research presented in this manuscript is primary research conducted at and supported by the National Research Council of Canada. All listed authors have reviewed the data presented here, attest to the validity of the data, and approve its publication.
- All data and materials presented in the manuscript will be made available under reasonable terms
- Authors declare no competing interests

## Contributions

- SMc, TN, AM, DB, KH, TS, RG and RW contributed to conceptualization, experimental design, analysis, and interpretation of results
- TN, AS, DB, SMa, CG, RP, and QZ contributed to technical development, performed experiments, and compiled data.
- SMc, TN, KH, and RW contributed to writing and editing the manuscript

## Supplementary Figure Legends

**Supplementary Figure 1:**
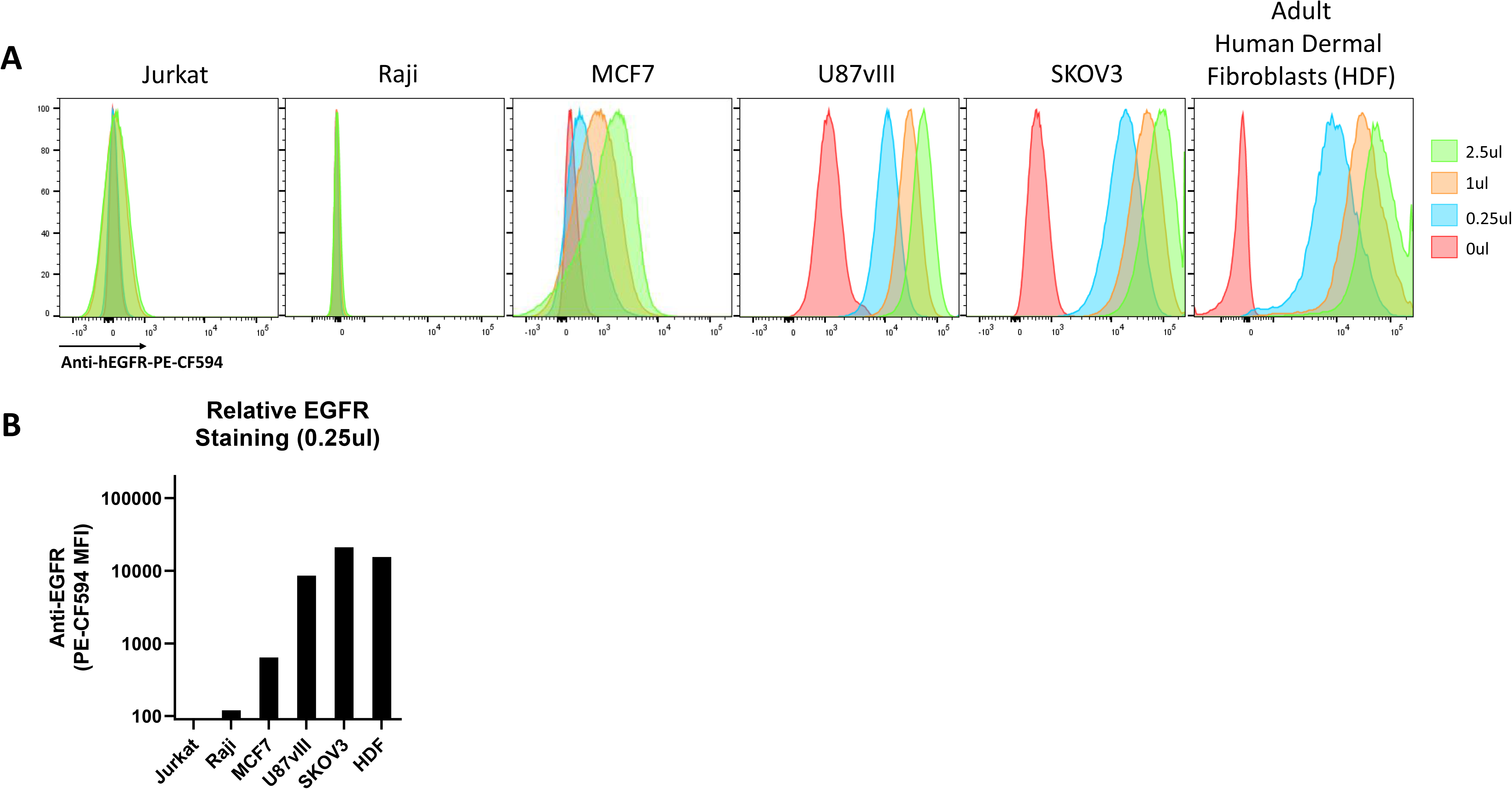
Measurement of surface EGFR expression on MCF7, U87vIII, and SKOV3 cells via flow cytometry. Jurkat, Raji, MCF7, U87vIII, SKOV3, and healthy donor derived primary human dermal fibroblast cells were stained with varying amounts of commercial PE-CF594 anti-human EGFR antibody and examined by flow cytometry as indicated in (A). (B) The mean fluorescence intensity of each cell type is shown. Results demonstrated lack of expression of EGFR in Jurkat and Raji, low expression in MCF7 cells, high expression in U87vIII, and very high expression for SKOV3and HDF cells. Histograms depict the fluorescence signal from a single experiment.

**Supplementary Figure 2:**
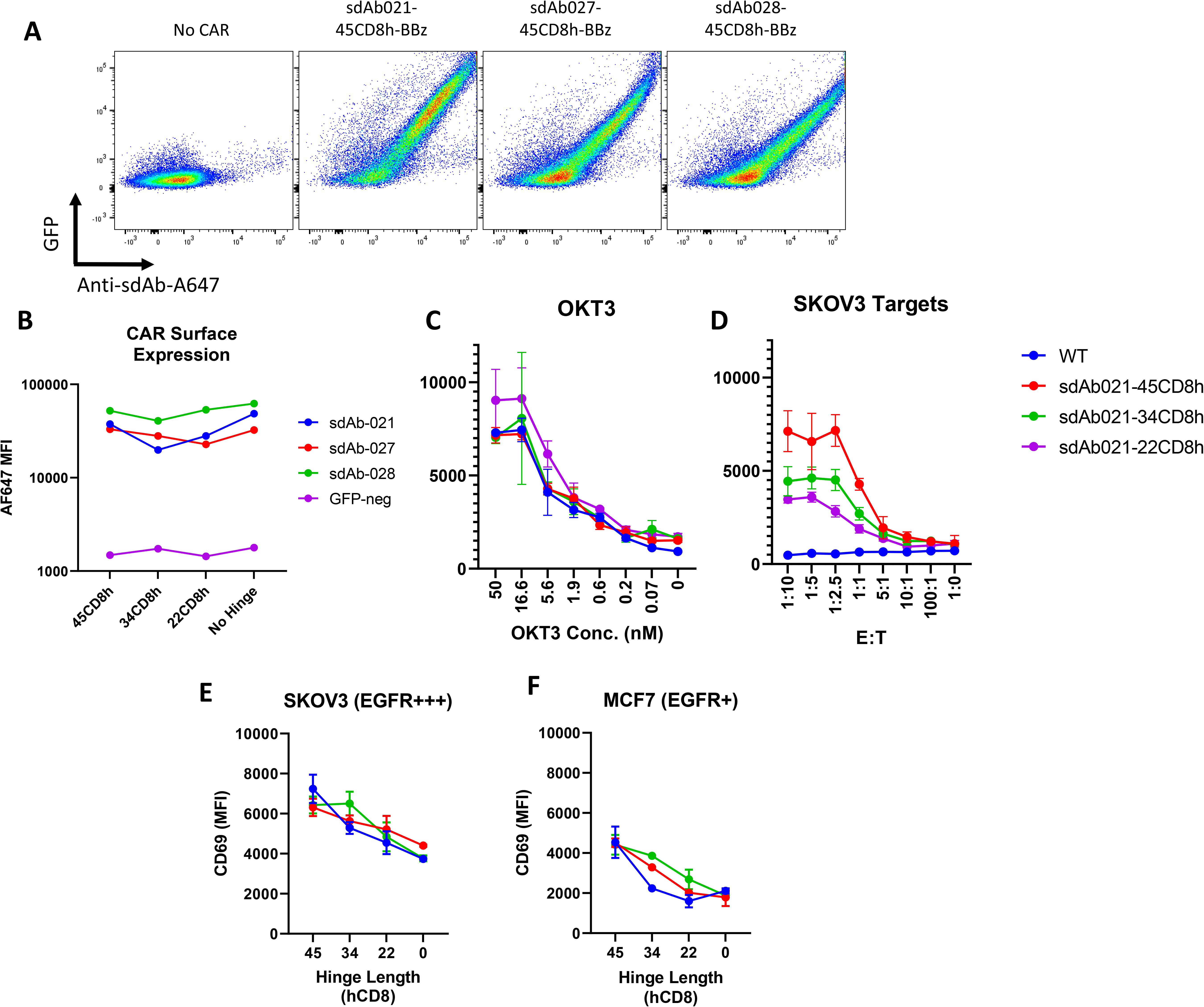
CAR Surface expression does not predict CAR response in EGFR-high and EGFR-low target cells. (A) Jurkat cells were electroporated with various EGFR-sdCAR plasmids were stained with an Alexa-Fluor647 labelled murine monoclonal anti-sdAb antibody to examine surface CAR expression. WT Jurkat cells (left panel) show no GFP and no binding to anti-sdAb, whereas all other EGFR-CARs show clear simultaneous GFP expression and binding to anti-sdAb. (B) Mean fluorescence intensity for AF647-anti-sdAb stained CAR-expressing Jurkat cells, gated on GFP expression is shown. (C) CAR-J CD69 expression of hinge modified EGFR sdCARs stimulated with plate bound OKT3 stimulation overnight are shown in comparison to (D) CAR-J cells stimulated with EGFR-high SKOV3 cells at varying dose. Results demonstrate modulation of antigen-specific response via the truncated hinge. (E) Data presented for CAR-J assay against EGFR-high SKOV3 cells as described in figure 2 were reorganized here to show the activation of EGFR sdCARs over a range of hinge lengths at an effector to target ratio of 1:10, varying from a full 45 amino acid human CD8 hinge to no hinge. (F) Data from CAR-J data against EGFR-low MCF7 cells organized via hinge length are shown.

**Supplementary Figure 3:**
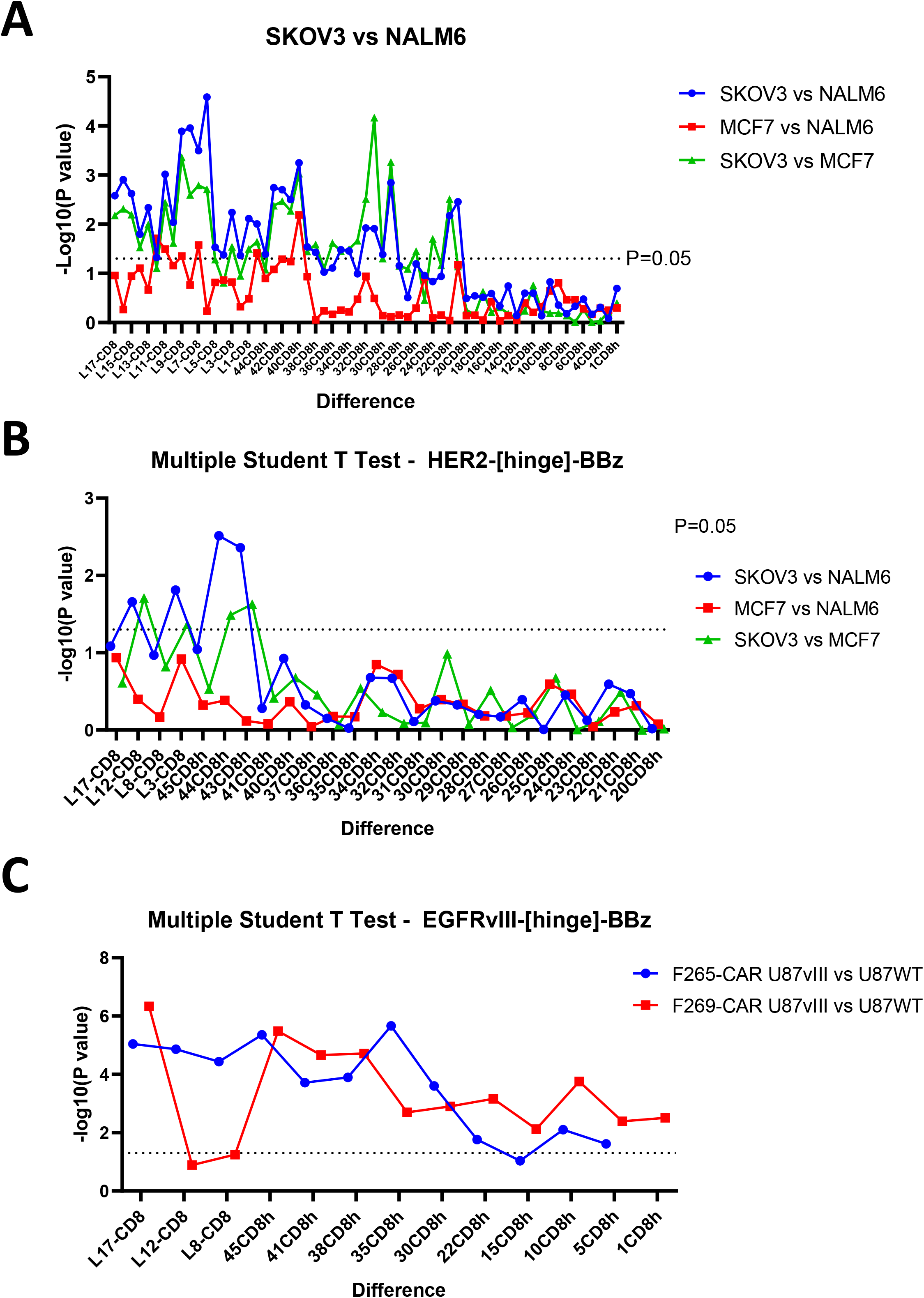
Statistical significance of divergence of CAR-T response against target antigen positive, low, or negative target cell lines. The mean the fold change in CD69 from 3 independent experiments as shown in Figure 3 were compared with similar construct responses to alternate target cells via the student T test (ie Fold change in CD69 in response to SKOV3 versus NALM6 for L17-CD8h-CAR). The negative log of the P value is shown with a dashed line at P=0.05. (A) Depicts the P value for statistical divergence of target response for various sdAb021-[hinge]-BBz constructs, (B) Depicts the P value for statistical divergence of target response for various HER2scFv-[hinge]-BBz constructs**, (C)** Depicts the P value for statistical divergence of target response for various EGFRvIIIscFv-[hinge]-BBz constructs.

**Supplementary Figure 4:**
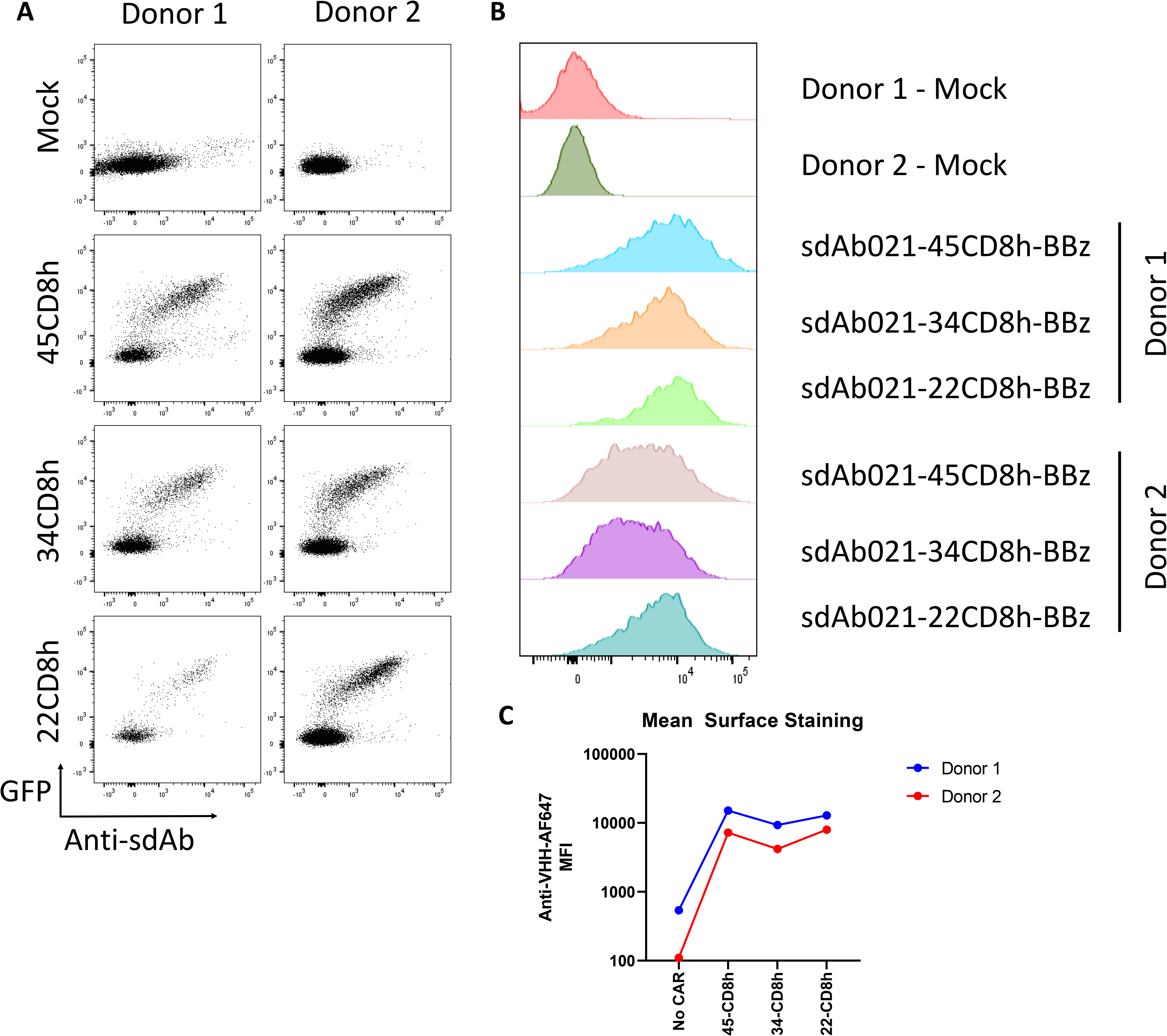
CAR Surface expression in primary CAR-T cells does not correlate with hinge length. (A) Primary T cells from two independent donors were purified, expanded, and transduced with CAR lentivirus as described in the methods section. CAR-T cells were stained with AlexaFluor647 anti-sdAb antibody and examined via flow cytometry to assess CAR surface expression. GFP/CAR+ T cells show a clear binding to the sdAb antibody, not seen in untransduced Mock T cells. (B,C) Histograms and mean fluorescence intensity for anti-sdAb stained CAR-T cells is shown, with no consistent effect of hinge truncation on CAR surface expression.

**Supplementary Figure 5:**
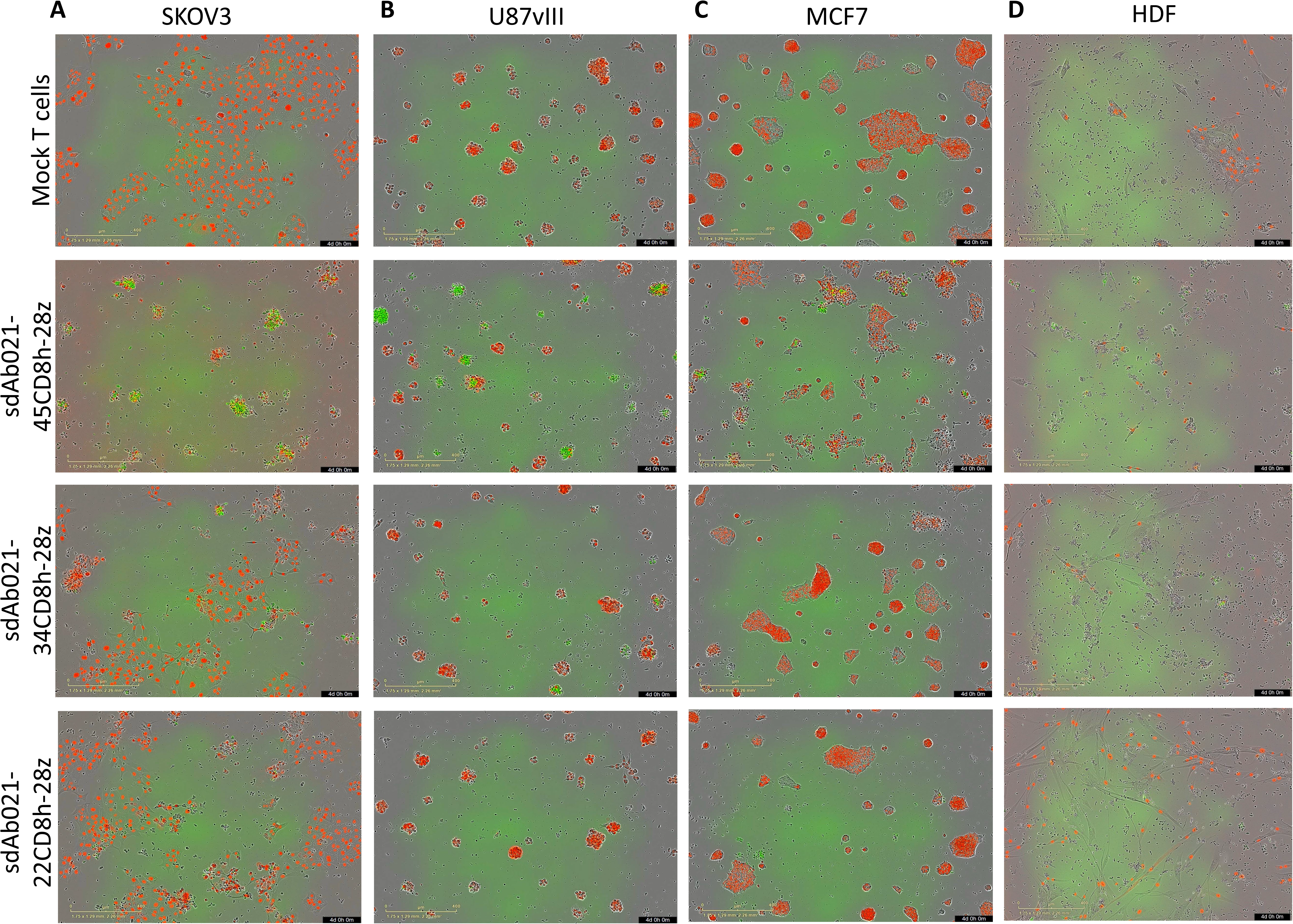
Hinge truncation results in reduced response to antigen-low and non-malignant EGFR+ target cells. (A and B) Primary sdCAR-T cells were generated from two independent blood donors as described in the text and placed at low density in a 96-well plate. Cells were then monitored for growth via live fluorescence microscopy over 7 days. Graphs display the total areas of GFP+ cells as enumerated through automated cell counting. (C) Representative images are provided for various sdCAR-T (GFP/green) or control mock T cells with target cell (mKate2/red) co-cultures 4 days after plating at an effector:target ratio of 5:1.

**Supplemental Figure 6:**
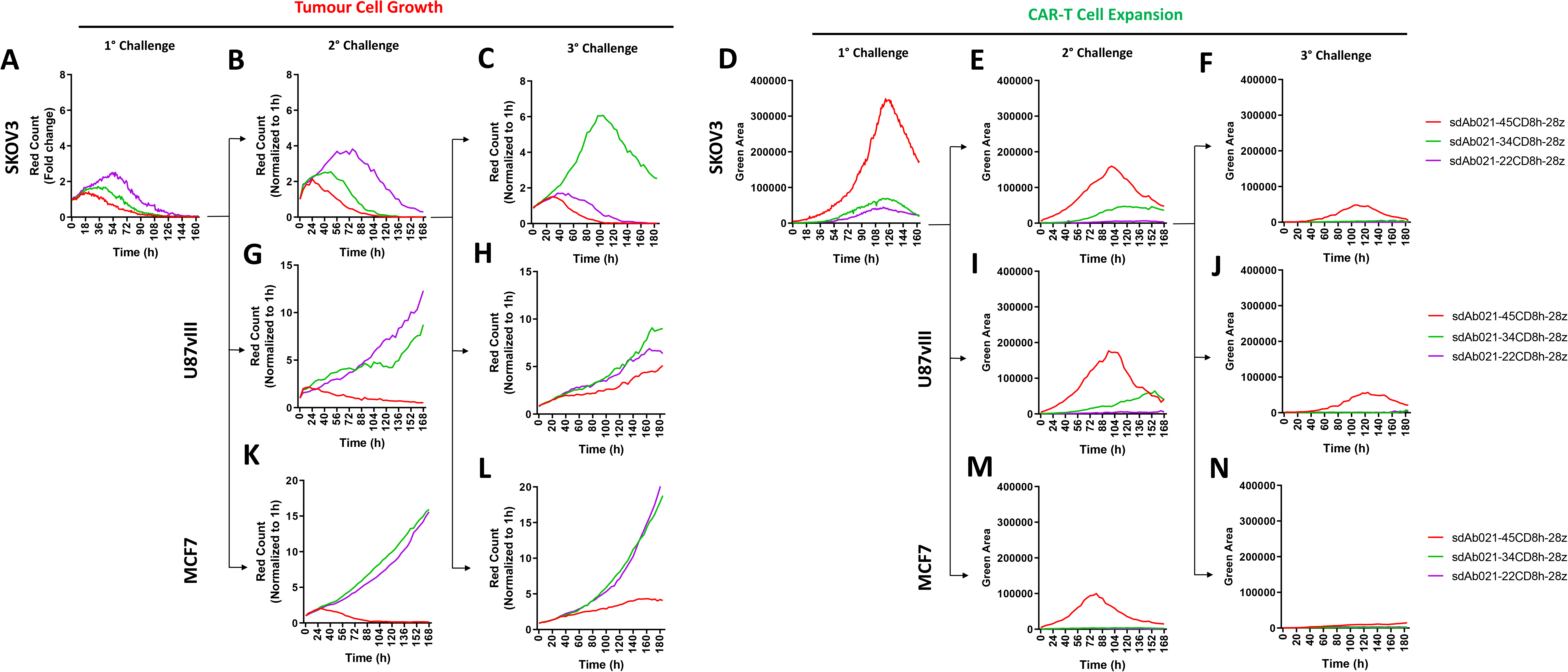
Hinge-truncated sdCAR-T cells maintain selectivity for EGFR overexpressing cells following re-challenge. Donor blood derived T cells were transduced with lentiviral particles encoding the EGFR-specific sdAb021 sdCAR with varying hinge domains. The resulting sdCAR-T cells were challenged via co-culture with EGFR-high SKOV3 cells and examined for (A) target cell expansion or (B) sdCAR-T cell expansion as described above. Challenged cells were then diluted 1/10 with fresh media and challenged with (B,E) EGFR-high SKOV3 cells, (G,I) EGFR-medium U87vIII cells, or (K, M) EGFR-low MCF7 cells. Similarly, cells challenged twice with SKOV3 cells were re-challenged with various target cells and examined for (C,H,L) target cell expansion and (F,J,N) CAR-T cell expansion. Each graph depicts automated cell counts or fluorescent areas from a single well from a single experiment.

**Supplementary Figure 7:**
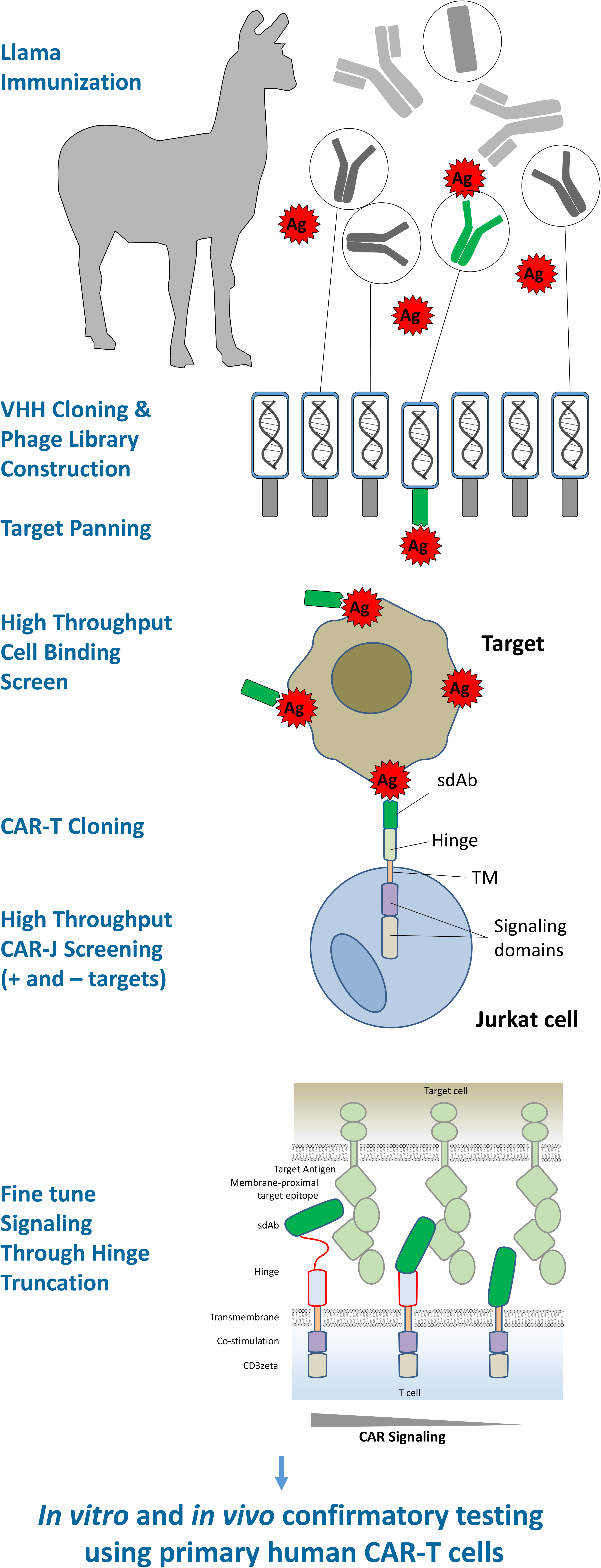
Workflow for discovery and optimization of novel tumor selective sdCAR constructs: Antigen preparation, llama immunization, phage library preparation and panning, cell binding, CAR cloning, high-throughput CAR screening, and fine-tuning of CAR signaling in Jurkat cells through hinge length modulation was performed for several novel EGFR sdCAR constructs. Subsequently, confirmatory experiments were performed to test lead sdCARs *in vitro* and *in vivo* using primary human CAR-T cells.

